# *Staphylococcus haemolyticus* is a reservoir of antibiotic resistance genes in the preterm infant gut

**DOI:** 10.1101/2025.01.25.634871

**Authors:** Lisa E. Lamberte, Elizabeth M. Darby, Raymond Kiu, Robert A. Moran, Antia Acuna-Gonzalez, Kathleen Sim, Alexander G. Shaw, J. Simon Kroll, Gusztav Belteki, Paul Clarke, Heather Felgate, Mark A. Webber, William Rowe, Lindsay J. Hall, Willem van Schaik

## Abstract

Among coagulase-negative staphylococci, *Staphylococcus haemolyticus* is a primary cause of bloodstream infections in preterm infants, with gut colonisation being recognised as a risk factor for subsequent infection. Through a re-analysis of a 16S rRNA gene sequencing dataset (n=497 preterm infants), we found that *S. haemolyticus* was abundant and prevalent in the gut in the first month of life. To better understand the diversity of *S. haemolyticus* among preterm infants, we generated genome sequences of *S. haemolyticus* strains (n=140), which were isolated from 44 stool samples of 22 preterm infants from four different hospitals in the United Kingdom. Core genome phylogenetic analyses, incorporating 126 publicly available *S. haemolyticus* genome sequences, showed that 85/140 (60.1%) of the isolates, from three different hospitals, formed a clonal group with 79/85 (92.9%) strains being assigned to Multi-Locus Sequence Type (ST) 49. Antibiotic resistance genes were highly prevalent in the genome sequences. Using logistic regression, we found a strong association between the presence of the gene *mecA* and phenotypic resistance to oxacillin (odds ratio [OR]: 158.00, p<0.0001), and the *aacA-aphD* gene and phenotypic resistance to gentamicin *aacA-aphD* (OR: 162.00, p<0.001). None of the strains from the preterm infant cohort had a complete Staphylococcal Cassette Chromosome *mec* (SCC*mec*) element. The *aacA-aphD* gene was associated with the transposon Tn*4001.* Using hybrid genome assemblies, we found it to be present on the chromosome (54.5% of strains) or on diverse plasmids (27.3%). Four strains (18.2%) had Tn*4001* copies on both plasmid and chromosome. Our data suggest the existence of a distinct sub-population of *S. haemolyticus* that has adapted to colonise the gut of preterm infants. Prevalent resistance to antibiotics is of clinical concern and the diversity of genetic contexts of *mecA* and Tn*4001* suggests widespread horizontal gene transfer and recombination in this species.

## Introduction

Preterm infants, defined as neonates born before 37 weeks of gestation, are a patient population that is particularly vulnerable to life-threatening complications. There are approximately 15 million preterm births globally, with an estimated 0.66 million deaths associated with preterm birth [1, 2]. The heightened vulnerability of preterm infants in neonatal intensive care units (NICUs) to nosocomial infections arises from their compromised immune defences, prolonged use of invasive medical tools, extended hospital stays, and concurrent medical complexities [3]. Late-onset sepsis (LOS) is a significant cause of morbidity and mortality among preterm infants, with coagulase-negative staphylococci (CoNS) as one of the leading causative agents of these infections [4–6]. Among CoNS, *Staphylococcus haemolyticus*, a skin commensal that commonly colonises the neonatal gut, is a prominent cause of LOS in preterm infants [7–9]. To treat neonatal sepsis, current UK guidelines recommend the use of a combination of antibiotics of multiple classes, primarily a β-lactam antibiotic (benzyl-penicillin or flucloxacillin) and the aminoglycoside gentamicin [10]. Among preterm infants hospitalised in UK NICUs between 2010 and 2017, 77% receive antibiotics at least once, with benzyl-penicillin and gentamicin being the most used antibiotics in these infants [11].

The emergence of multidrug-resistant strains among hospital-acquired pathogens in NICU settings poses a substantial challenge to treatment strategies and infant care [12, 13]. High-resolution genomic studies can provide a deep understanding of the intricate dynamics of these pathogens, shedding light on colonization patterns and the evolution of resistance mechanisms over time [14–18]. While *S. haemolyticus* is one of the most common CoNS causing human infections, it remains understudied and we thus lack an understanding of its diversity and potential for transmission in healthcare systems.

In this study, we employed high-throughput whole-genome sequencing to investigate the genomic diversity of *S. haemolyticus* that colonised the gut of preterm infants that were hospitalised in UK NICUs. We specifically focussed on its antibiotic resistance genes and the mobile genetic elements these genes are carried on, to provide insights into the threat posed by *S. haemolyticus* to critically ill preterm infants. Our findings can inform targeted strategies for better patient care and the management of neonatal infections, thus contributing to efforts that enhance clinical outcomes for vulnerable neonates.

## Materials and Methods

### 16S rRNA gene profiling of *S. haemolyticus* from a preterm infant stool sample dataset

We conducted 16S rRNA gene profiling to assess *S. haemolyticus* abundance in an existing dataset of 497 preterm infant stool samples from the Baby-Associated Microbiota of the Intestine (BAMBI) study (European Nucleotide Archive Accession Number: PRJEB31653) [19]. Sample details, including the proportion of reads assigned to *S. haemolyticus*, were extracted for analysis. Data visualization and figure generation were performed using RStudio (version 4.4.1), with the ggplot2 package (version 3.5.1).

### Isolation of *S. haemolyticus* from preterm infant stool samples

*S. haemolyticus* strains were isolated from preterm infant stool samples that were part of the Baby-Associated Microbiota of the Intestine (BAMBI) study [19] and the NeoM study [20]. A total of 140 isolates were collected from 44 stool samples of 22 infants. These stool samples originated from four hospitals in the East of England, specifically in Norwich; n = 28), two hospitals in London (London 1; n = 12; London 2; n = 3), and Cambridge (n = 1). Isolates were selectively cultured from stool samples using mannitol salt agar (Oxoid), on which *S. haemolyticus* forms pink colonies, and subsequently, on Tryptic Soy Agar (Thermo Scientific) + 5% sheep blood (Blood Agar), on which *S. haemolyticus* can be presumptively identified by beta-haemolysis. A total of 5 colonies per stool sample were collected and prepared for whole genome sequencing.

### Genomic DNA extraction and whole genome sequencing

Isolates that were sent for whole genome sequencing were grown from a single colony in Tryptic Soy Broth (Oxoid) for 18 h at 37°C. Genomic DNA for short-read (Illumina) sequencing was extracted using the Wizard Genomic DNA Purification Kit (Promega) according to the manufacturer’s instructions with a pretreatment of pelleted cells in 600 µl 50 mM EDTA with 1 mg/ml lysozyme and 1 mg/ml lysostaphin for 30 min at 37°C. Genomic DNA for long-read (Oxford Nanopore Technologies) sequencing was extracted using the Monarch HMW DNA Extraction Kit for Tissue (New England Biolabs) according to the manufacturer’s instructions with the addition of lysozyme and lysostaphin as above, prior to cell lysis. For short-read sequencing, DNA libraries were run at a final concentration of 1.5 pM, which included a 1% PhiX spike-in (PhiX Control v3 Illumina Catalogue FC-110-3001) on an Illumina Nextseq500 system using Mid Output Flowcells (NSQ® 500 Mid Output KT v2 (300 CYS) Illumina Catalogue FC-404-2003). For long-read sequencing, DNA libraries were prepared using the ligation sequencing kit SQK-LSK109 (Oxford Nanopore Technologies) and sequenced on the MinION using R9.4.1 flow cells, according to the manufacturer’s instructions.

### Genome Assembly

For short-read data, adapters were removed and trimmed using fastp (v.0.23.2) [21]. The short-read sequences were then assembled using SPAdes (v.3.14.1) with default parameters applied [22]. DNA assemblies were then annotated using Prokka (v.1.14.6) [23] and the sequence type (ST) was assigned using mlst (v.2.16.1) [24], using the PubMLST database [25].

The long-read data was basecalled using Guppy (v.0.1.0) [26]. The quality of the data was examined using NanoStat (v.1.6.0) [27]. Filtlong (v.0.2.1) [28] was used to filter out any reads shorter than 1kbp and exclude the worst 5% of the reads, using the option --keep_percent 95. Hybrid assemblies using the combination of long- and short-read data were generated using Unicycler (v.0.4.7) [29]. These assemblies were visualized in Bandage (v.0.8.1) [30]. In certain cases, Unicycler was unable to output a complete assembly resolution (i.e. generation of exclusively circular sequences). To address this, a long-read-first strategy was adopted. Specifically, Flye (v.2.9.1-b1780) [31, 32] was used to assemble the long read dataset, producing a .gfa file, which was integrated into the Unicycler pipeline to enhance the assembly process. SnapGene (www.snapgene.com) (v5.2.4) was used to visualise plasmids annotated by Prokka (v1.14.5) [23] that were generated from hybrid assemblies.

Assemblies were then searched for the presence of antibiotic resistance genes using ABRicate (v.1.0.1) [33] with the CARD database (v3.1.1) [34] and plasmid replicon sequences with the PlasmidFinder database (v2,0.1) [35, 36].

### Phylogenetic analysis

*S. haemolyticus* genomes from Cavanagh *et al.* [14] were added to the collection and were downloaded from the European Nucleotide Achive (PRJEB2705). Additionally, the genome sequence of the *S. haemolyticus* type strain NCTC11042 (Genbank: GCF_900458595.1) was included.

Annotated GFF files from Prokka (v.1.14.6) [23] were used as inputs for Roary (v.3.12.0) [37] to generate a core gene alignment using MAFFT at 95% sequence identity as determined by BLASTp, based on the concatenation of 1195 core genes. This core gene alignment was used to infer a maximum likelihood (ML) phylogeny using RAxML (v.1.1.0) with the GTR+G model with 100 bootstrap replicates [38]. Recombination events were removed using ClonalFrameML (v.1.12) [39]. snp-dists (v0.8.2) (https://github.com/tseemann/snp-dists) was used to quantify the number of SNPs in core genome alignments.The R package *fastbaps* [40] was used for conducting phylogenetic clustering analysis, enabling the identification of population structure. Detection of antibiotic resistance genes was performed using ABRicate (v.1.0.1) [33] on the genome assemblies with cut-offs of >80% coverage and >80% identity using the CARD [34] and VFDB [41] databases. Phylogenetic trees were visualized and annotated using iTOL [42].

### Identification of SCC*mec*-related clusters

The initial SCC*mec* typing of *S. haemolyticus* assemblies was performed using SCC*mec*Finder [43], but complete SCC*mec*s were not detected with this tool. To further assess the presence of SCC*mec*-encoded elements, a reference dataset comprising 112 SCC*mec* elements, representing SCC*mec* types I to XIV, was compiled (Table S1). These elements were sourced from diverse reference strains in accordance with the classification system established by the International Working Group on the Staphylococcal Cassette Chromosome (IWG-SCC, 2022) and retrieved from the NCBI GenBank database [44]. Each *S. haemolyticus* assembly from our study collection was compared against this reference database using BLASTn [45]. SCC*mec* elements were identified based on criteria of >80% sequence coverage and >80% nucleotide identity.

### Antibiotic susceptibility testing

Antibiotic susceptibility testing was performed on a total of 58 *S. haemolyticus* isolates from the BAMBI collection. Susceptibility to gentamicin and oxacillin was determined utilising the broth microdilution method, as described [46], and the results were interpreted according to EUCAST breakpoints [47]. The reference strain *S. haemolyticus* NCTC11042 was used as a control. The minimum inhibitory concentration (MIC) was determined as the mode of three biological replicates.

Furthermore, the analysis was expanded by incorporating antibiotic susceptibility data from an additional 123 *S. haemolyticus* strains from a study by Cavanagh and colleagues [14], resulting in a total of 181 strains for which we had phenotypic antibiotic susceptibility data.

### Statistical analyses

All statistical analyses were performed using R version 4.2.2. We employed unconditional multivariable logistic regression to investigate the association between the presence of genes and phenotypic resistance. The dataset, comprising 181 isolates, was organised into a binary outcome variable denoting phenotypic resistance and predictor variables representing the presence or absence of genetic elements. The logistic regression model was fitted using the glm function in R, and the resulting model summary was examined for coefficient estimates, odds ratios, and 95% confidence intervals. The statistical significance of each genetic element’s association with phenotypic resistance was assessed based on the p-values, with a significance level set at p <0.05.

### Data Availability

Raw sequencing reads generated in this study have been submitted to Genbank under the BioProject PRJNA1105567.

## Results

### *S. haemolyticus* is abundant and highly prevalent in stool samples of preterm infants hospitalised in the NICU

To determine the prevalence of *S. haemolyticus* in the preterm infant gut microbiota, we re-analyzed 16S rRNA gene sequencing data generated on stool samples (n=497), from preterm infants (n=192) enrolled in the Baby-Associated MicroBiota of the Intestine (BAMBI). The stool samples were collected at four timepoints (0 – 9, 10 – 29, 30 – 49, and 50 – 99 days from birth) [19]. This re-analysis showed that *S. haemolyticus* was initially abundant in the first 10 days after birth (Fig. 1). While the abundance of *S. haemolyticus* decreased as the infants aged, *S. haemolyticus* was still detectable in some infants at later time points (30-70 days of age). Specifically, *S. haemolyticus* was detected in 22.2% of samples at time point 0 – 9 days, 29.5% of samples at 10 – 29 days, 11.7% of samples at 30 – 49 days, and 6.4% of samples at 50 – 99 days from birth.

**Figure 1.**
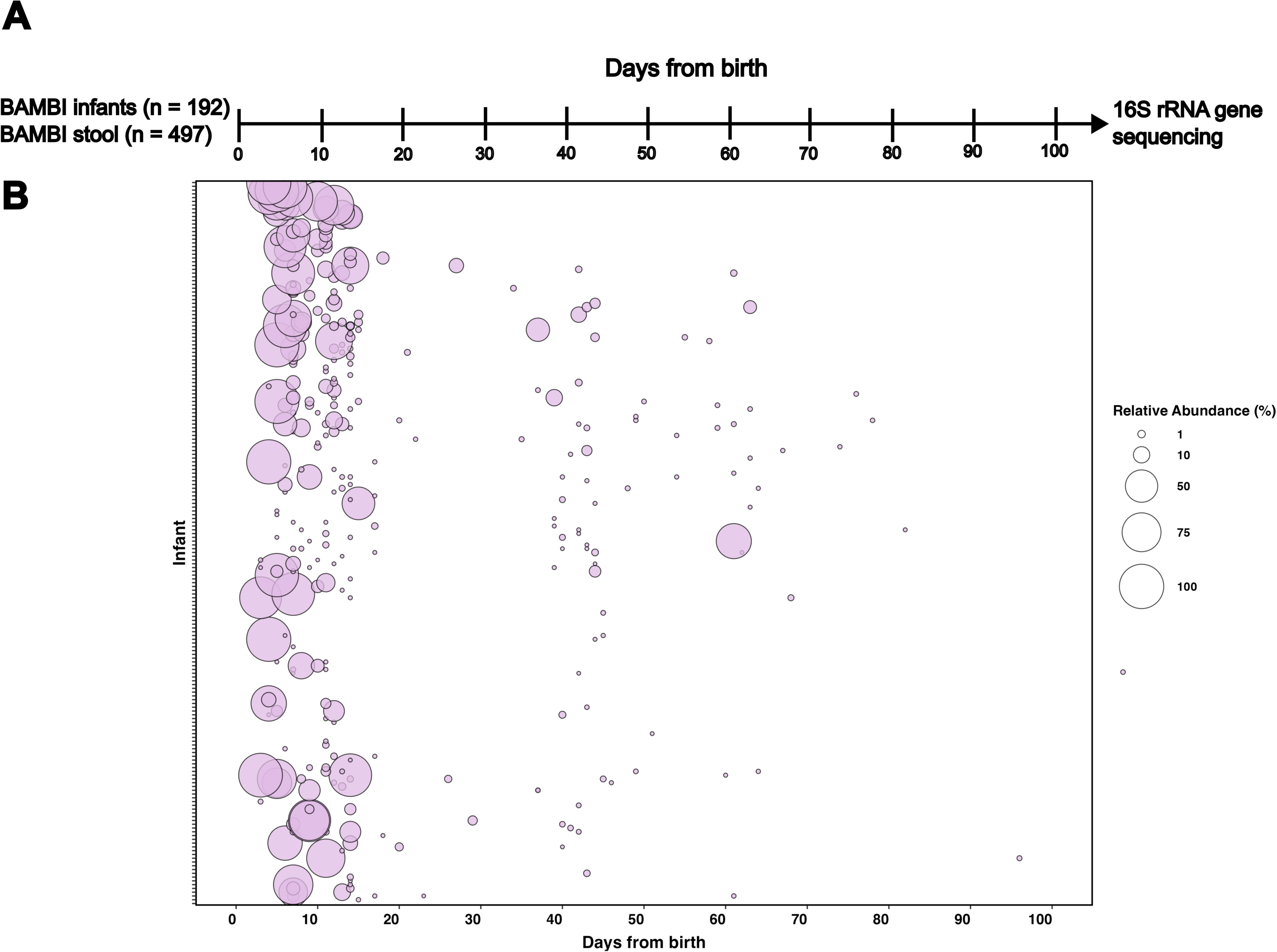
Detection of *Staphylococcus haemolyticus* in the preterm infant gut microbiota. A total of 497 stool samples were collected from 192 patients and were subjected to 16S rRNA gene sequencing *[19]*, with data. Bubble plots show the relative abundance of *Staphylococcus haemolyticus* detected in the collection of stool samples (n=497).

### *S. haemolyticus* isolates detected in majority of preterm infant stool samples belong to a clonal population

To assess the genetic diversity of *Staphylococcus haemolyticus* in stool samples from the BAMBI preterm infant cohort, we isolated presumptive *S. haemolyticus* strains from available stool specimens (n = 46) obtained from preterm infants (n = 22) (Fig. S1). Whole genome sequencing confirmed the identity of 140 strains from 44 samples as *S. haemolyticus*, while the remaining 10 strains (from 2 samples) were identified as *Staphylococcus epidermidis*. After genome sequencing of the *S. haemolyticus* isolates, we constructed a core genome phylogeny (Fig. 2) to study the relatedness of *S. haemolyticus* isolated from stool samples of preterm infants across four UK hospitals. This phylogeny was based on the 140 whole genome sequences from strains isolated here and 126 publicly available *S. haemolyticus* genomes. Fastbaps clustering analysis divided the 266 *S. haemolyticus* isolates into 22 clusters. The largest cluster, Cluster 1, consisted of 85/266 (32.0%) isolates, all coming from this study’s preterm infant cohort with isolates from three UK hospitals. Individual isolates differed between 0 and 369 core genome SNPs (cgSNPs) with an average cgSNP difference between two Cluster 1 genomes of 64.6 SNPs. Out of 916 instances in which cluster 1 strain pairs were closely related to each other (≤10 core genome SNPs [cgSNPs]), 256 (27.9%) were found to be shared between different hospitals, suggesting that Cluster 1 represents a geographically dispersed, but host-adapted population of *S. haemolyticus*. Within infants, we also noted significant variability. If an infant was only colonised by Cluster 1 strains, the number of cgSNPs between strains isolated from the same stool sample ranged between 0 and 251 cgSNPs. Among the 44 stool samples from which *S. haemolyticus* was isolated, three (8.3%) returned strains from different clusters.

**Figure 2.**
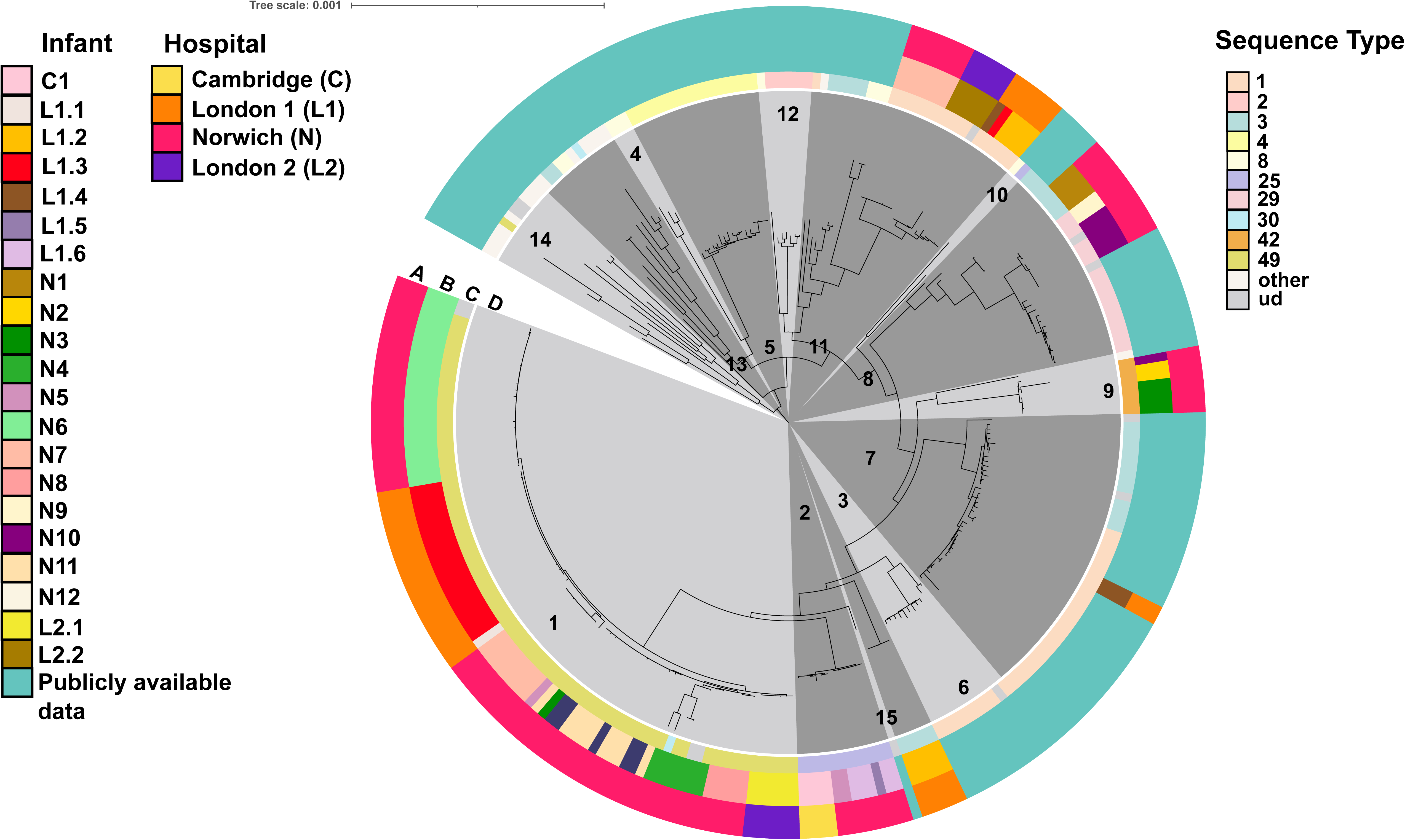
Midpoint-rooted phylogenetic tree illustrating core genome comparisons among strains sequenced in this study and publicly available strains (n = 266). The concentric rings display metadata related to stool isolates collected from preterm infants at various hospitals across the UK. Ring A indicates the hospital of origin for each isolate. Ring B denotes individual infants from whom the strains were isolated. Ring C shows multilocus sequence typing (MLST) designations for each strain. The central ring (Ring D) uses alternating dark and light grey blocks to represent Fastbaps clusters, numbered for reference. Sequence types (STs) with three or more isolates are labeled individually in the Sequence Type legend, while STs with fewer than three isolates are grouped under “Other.” Strains for which sequence types could not be determined are labeled as “ud” (undetermined).

In the *S. haemolyticus* genome sequences, we identified multiple antibiotic resistance genes (ARGs) that are predicted to confer resistance to 13 antibiotic classes (Fig. 3). The most commonly detected ARGs in our preterm infant cohort were the regulatory gene *mgrA* (140/140; 100%) the aminoglycoside resistance gene *aacA-aphD,* and the β-lactam resistance genes *blaZ* and *mecA* (all 139/140; 99.3%). The gene *qacA* (132/140; 94.3%), an efflux pump associated with tolerance towards disinfectants, was also nearly ubiquitous among the isolates from the preterm infant cohort.

**Figure 3.**
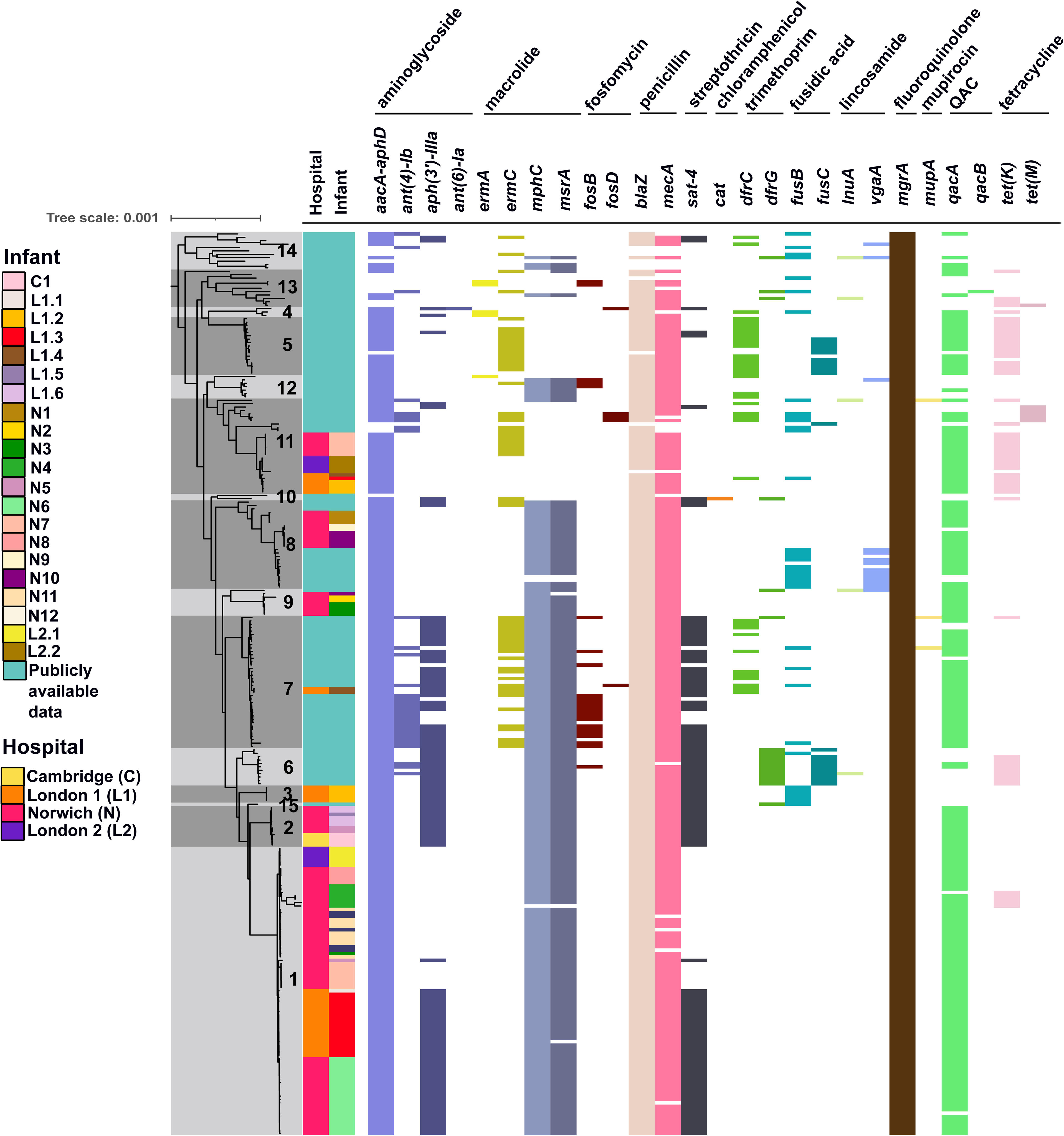
Antibiotic resistance gene (ARG) profiles in *Staphylococcus haemolyticus* isolates. This figure displays the presence or absence of specific ARGs and their associated antibiotic classes across isolates from preterm infants, grouped by their hospitals of origin. A core genome phylogenetic tree, with Fastbaps clusters indicated, is shown on the left, together with information on the hospitals each infant was hospitalized in. Abbreviations: QAC – quaternary ammonium compounds.

We then investigated the association between genotypic and phenotypic resistance to the antibiotics oxacillin (a beta-lactam) and gentamicin (an aminoglycoside). Out of the 176 *S. haemolyticus* isolates tested for oxacillin resistance, 158 isolates harbouring the *mecA* gene exhibited phenotypic resistance to oxacillin (158/176; 89.8%), 1 isolate harbouring the *mecA* gene exhibited phenotypic sensitivity to oxacillin (1/176; 0.6%), 6 isolates lacking the *mecA* gene exhibited phenotypic resistance to oxacillin (6/176; 3.4%), while 11 isolates lacking the *mecA* gene exhibited phenotypic sensitivity to oxacillin (11/176; 6.3%). An unconditional multivariable logistic regression was employed to assess the association between the presence of genes to phenotypic outcome. The logistic regression model revealed a strong association between the presence of the gene *mecA* and resistance to oxacillin (odds ratio [OR]: 158.00, 95% confidence interval [CI]: 134.63 – 183.92, p<0.0001), in contrast to the presence of the other beta-lactam resistance gene, *blaZ* (OR: 0.97, 95% CI: −0.78 – 1.22, p = 0.82) (Fig. 4).

**Figure 4.**
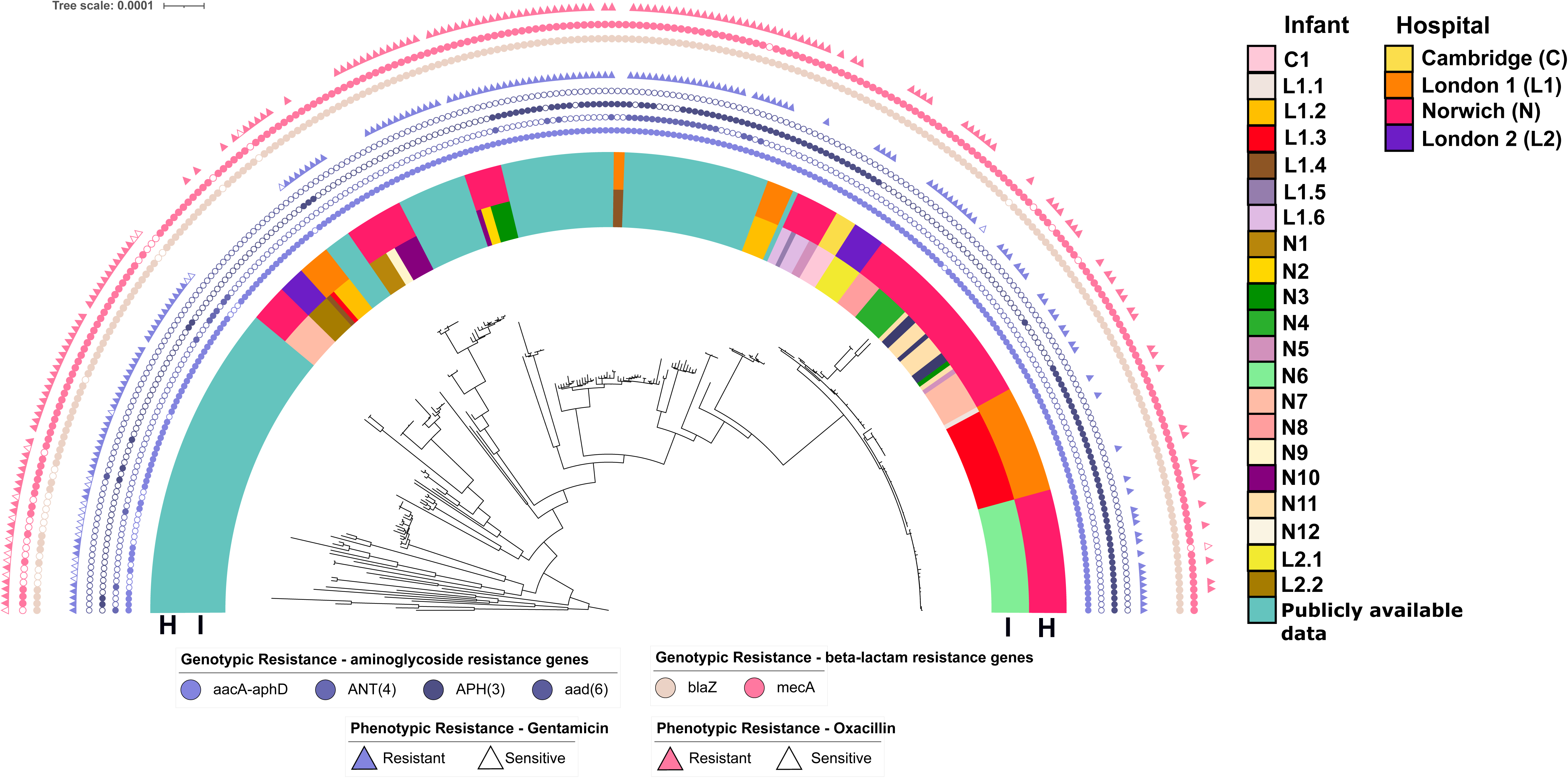
Presence of aminoglycoside and beta-lactam resistance genes and outcome of antibiotic susceptibility testing. Coloured purple circles represent the presence of aminoglycoside resistance genes *aacA-aphD, ant(4’)-Ib*, *aph(3’)-IIIa*, and *ant(6)-Ia*. Coloured pink circles represent the presence of beta-lactam resistance genes *blaZ* and *mecA*. Filled circles indicate that the gene is present, open circles indicate that the gene is absent. Filled triangles indicate phenotypic resistance to an antibiotic, open triangles indicate phenotypic sensitivity to antibiotics. The half-circle labeled “I” represents individual infants, and the half-circle labelled “H” indicates the hospitals from which these infants’ isolates were obtained, with the same colour coding as in Figures 2 and 3. The midpoint-rooted phylogenetic tree is based on a core genome alignment of *S. haemolyticus* genomes.

Of the 181 *S. haemolyticus* isolates tested for gentamicin resistance, 162 isolates encoding the *aacA-aphD* gene exhibited phenotypic resistance to gentamicin (162/181; 89.5%), whereas only 2 isolates encoding the *aacA-aphD* gene exhibited phenotypic sensitivity to gentamicin (2/181; 1.1%). Additionally, only 2 isolates lacking the *aacA-aphD* gene exhibited phenotypic resistance to gentamicin (2/181; 1.1%), while 15 isolates lacking the *aacA-aphD* gene exhibited phenotypic sensitivity to gentamicin (15/181; 8.3%). We found a significant positive association for the presence of *aacA-aphD* and phenotypic resistance to gentamicin (OR: 162.00, 95% CI: 138.31 – 188.24, p < 0.001), in contrast to the presence of other aminoglycoside genes (*ant(4’)-Ib* (OR = 0.18, 95% CI: 0.12 - 0.26, p<0.001), *aph(3’)-IIIa* (OR = 0.46, 95% CI: 0.35 – 0.60; p<0.001), and *ant(6)-Ia* (OR = 0.0061, 95% CI = −3.51 x 10^-4^ – 0.027; p<0.001) that were identified in our dataset.

### SCC*mec* elements are differentially distributed in *S. haemolyticus* isolates from the preterm infant cohort

The distribution of SCC*mec* core elements *orfX* (*rlmH*), *mecA*, *mecR*, *mecI*, IS*431*, *ccrA*, *ccrB*, and *ccrC* [48] was investigated in our collection of *S. haemolyticus* isolates. Hierarchical clustering of observed SCC*mec* elements per isolate revealed distinct patterns of SCC*mec* components between isolates from our preterm infant cohort and publicly available data (Fig. 5). A majority of *S. haemolyticus* isolates from our preterm infant BAMBI cohort (103/140; 73.6%) carried *orfX* (*rlmH*), *mecA*, and IS*431*. A smaller proportion of the isolates carried additional SCC*mec* elements, 19.3% (27/140) of isolates carried *orfX* (*rlmH*), *mecA*, IS*431,* and *ccrC* (27/140; 19.3%), 6.4% (9/140) of isolates had *orfX* (*rlmH*), *mecA*, IS*431, ccrA* and *ccrB* (9/140; 6.4%). Only a single isolate carried *orfX* (*rlmH*), *mecA*, IS*431, ccrA*, *ccrB* and *ccrC* (Fig. S2). None of the isolates from our study had a complete SCC*mec* element as the *mecR* gene was missing from these isolates. SCC*mec* elements were also incomplete in the publicly available genomes we included in this study. Utilising hybrid-assembled genomes, we examined the genetic context of the *mecA* regions within our *S. haemolyticus* preterm infant cohort and identified seven different genetic contexts (Fig. S2).

**Figure 5.**
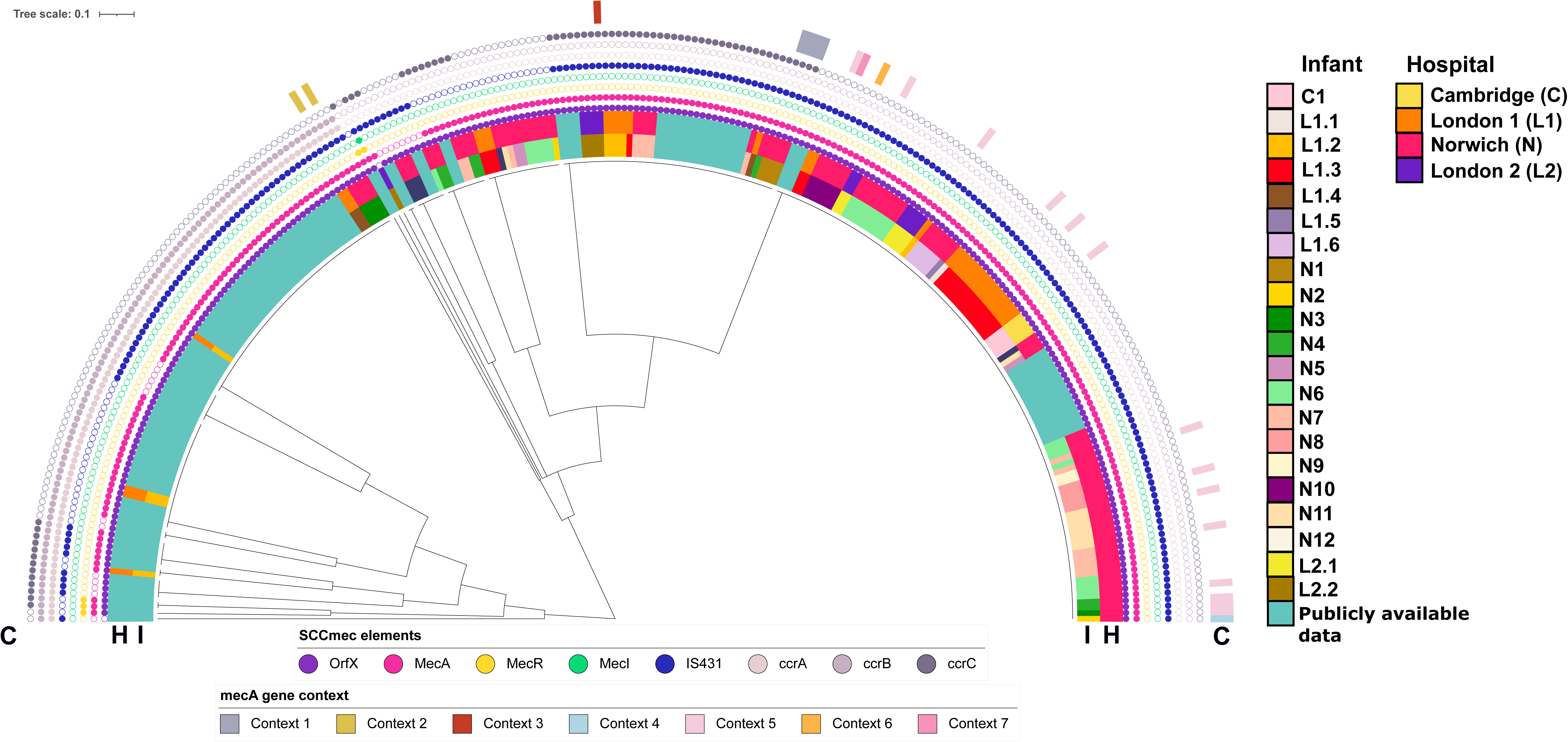
Genetic context of the *mecA* regions found in *S. haemolyticus* isolates from the preterm infant cohort. The tree visualises a hierarchical clustering of SCC*mec* core elements found in strains sequenced for this study, in addition to publicly available *S. haemolyticus* genomes. Coloured circles represent the presence (filled circle) or absence (open circle) of SCC*mec* core elements *orfX, mecA, mecR, mecI,* IS*431*, *ccrA, ccrB, and ccrC.* The half-circle labelled “C” represents the *mecA* genetic context (Fig. S2), the half-circle labelled “I” represents individual infants, and the half-circle labeled “H” indicates the hospitals from which these infants’ isolates were obtained, with the same colour coding as in Figs. 1 and 2.

Plasmid- and chromosome-mediated gentamicin resistance in *S. haemolyticus* isolates is encoded by a transposon, Tn*4001*.

We next examined the genetic context of gentamicin resistance gene *aacA-aphD* in the *S. haemolyticus* isolates. In the genomes we generated for this study, this gentamicin resistance gene is always associated with a transposon, Tn*4001*, which has first been described in *S. aureus* [49]. To conclusively map the genetic context of Tn4001, we generated hybrid assemblies for 22 strains and found that Tn*4001* was present on the chromosome in 12/22 (54.5%) of strains, and on a plasmid in 6/22 (27.3%) strains. In 4/22 (18.2%) strains Tn*4001* was present on both a plasmid and the chromosome (Figure 6).

**Figure 6.**
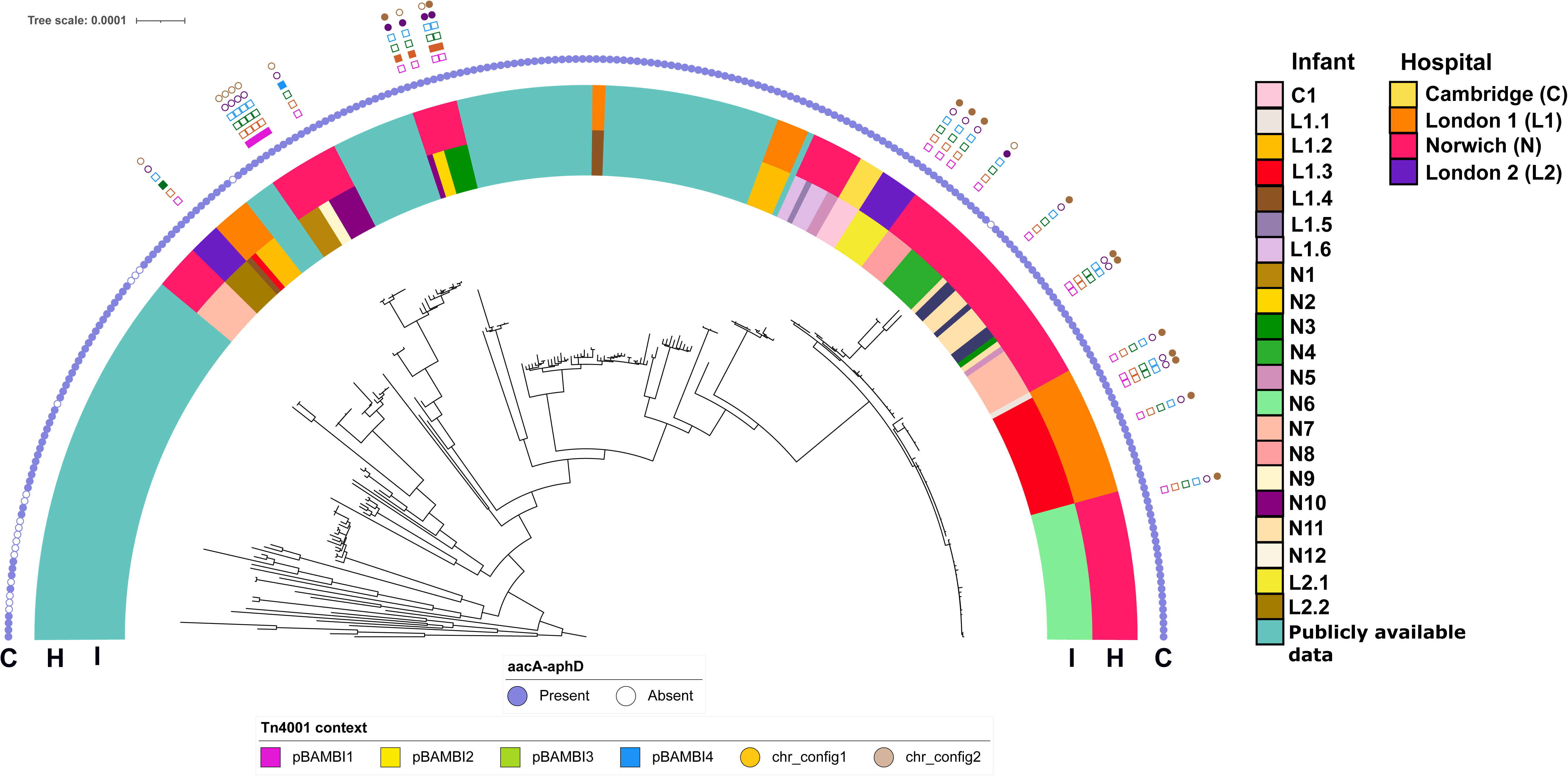
Genetic context of the Tn*4001* region found in *S. haemolyticus* isolates from the preterm infant cohort. Coloured purple circles represent the presence (filled circle) or absence (open circle) of *aacA-aphD* gene. Different genetic contexts of Tn*4001* are indicated: squares represent the configuration of Tn*4001* in plasmids (pBAMBI1 – pBAMBI4) and circles represent the configuration of Tn*4001* in the chromosome (chr1_config1 and chr_congfig 2) (Fig. S2). The half-circle labelled “C” represents the *mecA* genetic context (Fig. S2), the half-circle labelled “I” represents individual infants, and the half-circle labeled “H” indicates the hospitals from which these infants’ isolates were obtained and these use the same colours as in Fig. 1 and 2. The midpoint-rooted phylogenetic tree is based on a core genome alignment of *S. haemolyticus* genomes.

The transposon was found in four distinct plasmids, which we have named pBAMBI1, pBAMBI2, pBAMBI3, and pBAMBI4 (Fig. S3). In addition to Tn*4001*, all four of these plasmids also carry the *blaZRI* gene cluster, of which the first gene encodes an extracellular beta-lactamase that is widespread in *Staphylococcus*, and the *qacRA* genes [50], which contribute to resistance to quaternary ammonium compounds (QACs). We found two distinct configurations of the Tn*4001* chromosomal integration among the *S. haemolyticus* genomes (Fig. S3), which we termed chr_configs 1 and 2 (Figure 6).

## DISCUSSION

CoNS are currently understudied, compared to *S. aureus*, even though they are a frequent cause of infection in neonates [51]. There exists considerable diversity among CoNS and among this diverse group of species, several are significant opportunistic pathogens. After *S. epidermidis*, *S. haemolyticus* is among the CoNS most commonly causing bloodstream infections [52]. Preterm infants are at high risk of LOS caused by nosocomial CoNS strains, of which *S. haemolyticus* is one of the most prominent causes of infection. Gut colonisation by *S. haemolyticus* may be a risk factor for subsequent systemic infections in preterm infants [53]. Prevalence of *S. haemolyticus* in the preterm infant gut is particularly high in the first few weeks post-partum [54, 55]. In this study, we first re-analysed 16S rRNA gene sequencing data of stool samples of preterm infants, revealing a distinct temporal pattern, with an initial high abundance of *S. haemolyticus* in the first ten days after birth, followed by a gradual decline as infants mature. Notably, some infants continue to harbour high levels of *S. haemolyticus* even at later stages, suggesting a persistent colonisation within the NICU environment. We then employed high-throughput whole-genome sequencing to investigate the occurrence of *S. haemolyticus* within the preterm infant gut during their initial days of life

The core genome phylogenetic analysis revealed a diverse population of *S. haemolyticus* among the isolates obtained from different UK hospitals. Among this population, Cluster 1 was very prominent among preterm infant isolates. Additionally, Cluster 1 strains were identified in three different hospitals, demonstrating that Cluster 1 may represent a disseminated clonal population that has adapted to the gut colonisation of the preterm infant gut, similar to what was proposed for a sub-population of *Staphylococcus capitis* [56]. We currently cannot determine how highly similar *S. haemolyticus* strains have emerged in multiple hospitals. It is likely that transfer of patients or healthcare workers between hospitals, particularly between the two geographically close hospitals in London which are in the same National Health Service Trust, has led to the dissemination of Cluster 1 strains, similar to how methicillin-resistant *S. aureus* has spread among hospitals in the UK [57–59].

Our whole genome sequence analysis identified the presence of multiple antibiotic resistance genes (ARGs) in *S. haemolyticus* from preterm infants, with particularly high prevalence of genes conferring resistance to beta-lactams and aminoglycosides. The detection of these ARGs, particularly *aacA-aphD* and *mecA*, is intriguing, as the antibiotics gentamicin and flucloxacillin are frequently the antibiotics of choice for babies with late-onset neonatal infection who are already in a neonatal unit [10, 60]. Interestingly, using comparative genomics of nosocomial and commensal *S. haemolyticus* isolates, Pain and colleagues [61] have proposed that the presence of both these genes (i.e. *aacA-aphD* and *mecA*) are primary indicators of hospital adaptation and pathogenicity, whereas absence of these genes indicate a commensal isolate, and our data are in line with these observations [61]. The near ubiquitous spread of beta-lactam and gentamicin resistance genes in *S. haemolyticus* isolates of preterm infants raises important questions about the selective pressures, and potentially the efficacy of treatment of *S. haemolyticus* infections in preterm infants with these antibiotic classes. Indeed, bloodstream infections caused by *S. haemolyticus* may be associated with higher morbidity in neonates, compared to the more commonly encountered species *S. epidermidis* [62]. It is possible that a combination of multidrug-resistance and virulence, which is at least partially due to the production of a range of toxins and haemolysins, makes *S. haemolyticus* a particularly problematic CoNS in severely immunocompromised patient groups [63–65].

Our study revealed distinct and diverse genetic contexts of *mecA* regions and *ccr* gene complexes in *S. haemolyticus* isolates from preterm infants. While *S. haemolyticus* does not appear to be the direct source of SCC*mec* elements that contribute to methicillin resistance in *S. aureus*, the wide variety of genetic contexts of *mecA* in *S. haemolyticus* suggests significant rearrangements of this important resistance determinant within the species. We found that the gene *aacA-aphD*, responsible for gentamicin resistance, is encoded within a transposon, Tn*4001* [49]. This transposon was found in various genetic contexts, including on plasmids and in chromosomes, or in multiple copies distributed among both replicons, further highlighting the potential for Tn*4001*’s mobility within *S. haemolyticus*. Our findings on significant within-host diversity of *S. haemolyticus* can inform future studies into this microbe as it underscores the importance of collecting multiple isolates from a single sample to capture the diversity of *S. haemolyticus* in its environment. *S. haemolyticus* remains relatively understudied and future research could focus on generating a deeper understanding of the mechanisms by which *S. haemolyticus* can acquire, and potentially further disseminate, mobile genetic elements that carry antibiotic resistance genes.

## Author Contributions

L.E.L isolated strains and performed experiments. L.E.L, E.M.D, R.K, R.A.M., A.A.G., H.F. and W.R. performed bioinformatic analyses with advice from M.A.W. K.S., A.G.S., J.S.K, G.B., P.C supervised sample collection at their sites. L.E.L., L.J.H., and W.v.S. conceived the study and wrote the manuscript. All authors reviewed and approved the final manuscript.

## Ethical Statement

Faecal collection from Norfolk and Norwich University Hospital (NNUH) and Addenbrooke’s Hospital (BAMBI study) was approved by the Faculty of Medical and Health Sciences Ethics Committee at the UEA and followed protocols laid out by the UEA Biorepository (licence no. 11208). Faecal collection from Imperial Healthcare NICUs was approved by West London Research Ethics Committee (REC) under the REC approval reference no. 10/H0711/39.

## Supplemental figure legends

**Fig. S1. Overview of samples.** Children are indicated by unique identifiers. Closed circles denote samples from which *S. haemolyticus* was isolated. Open circles denote samples that did not yield *S. haemolyticus* upon culture. Filled squares represent individual infants.

**Fig. S2. Genetic context of SCC*mec* components in *S. haemolyticus* genomes and comparison with *S. aureus* prototypes.** Scaled linear diagrams of 7 different contexts of the *mecA* region. SCC*mec* elements (*orfX, mecA, ccrA, ccrB,* and *ccrC*), as well as genes of high similarity, are indicated as coloured arrows. ORFs and genes of low similarity are depicted as light green arrows. Sequences of SCC*mec* types V (5C2), V(5C2&5), VIII (4A), and X (7C1) are from Genbank accessions AB512767.1, AB121219.1, FJ390057.1, and AB505630.1, respectively. Diagrams are visualized using Clinker.

**Fig. S3. Genetic context of Tn*4001* in *S. haemolyticus* genomes.** (A) Circular map of plasmids pBAMBI1, pBAMBI2, pBAMBI3, and pBAMBI4. Plasmid sequence is shown as a black line, with arrows inside representing genes (coloured arrows) and genes encoding hypothetical proteins (light green). (B) Scaled linear diagrams of 6 different genetic contexts of the Tn*4001* region. Genes of high similarity, as well as genes unique to the *S. haemolyticus* isolates from the cohort are indicated as coloured arrows. ORFs encoding hypothetical proteins and genes of low similarity are depicted as light green arrows. Sequences of plasmid references are from Genbank Accessions: *Staphylococcus epidermidis* RP62A plasmid pSERP (CP000028.1), *Staphylococcus epidermidis* plasmid SAP016A (GQ900381.1), *Staphylococcus epidermidis* strain TMDU-265 plasmid p1 (CP093182.1), *Staphylococcus aureus plasmid* pSK1 (GU565967.1), Staphylococcus epidermidis strain 107.2cured plasmid pAQZ2 (MK046688.1), and *Staphylococcus haemolyticus* strain 12b plasmid pSH_12b_1 (CP071506.1). Diagrams are visualized using Clinker.

