## Supplementary figures and images for "*Staphylococcus haemolyticus* is a reservoir of antibiotic resistance genes in the preterm infant gut"

### Fig. S1

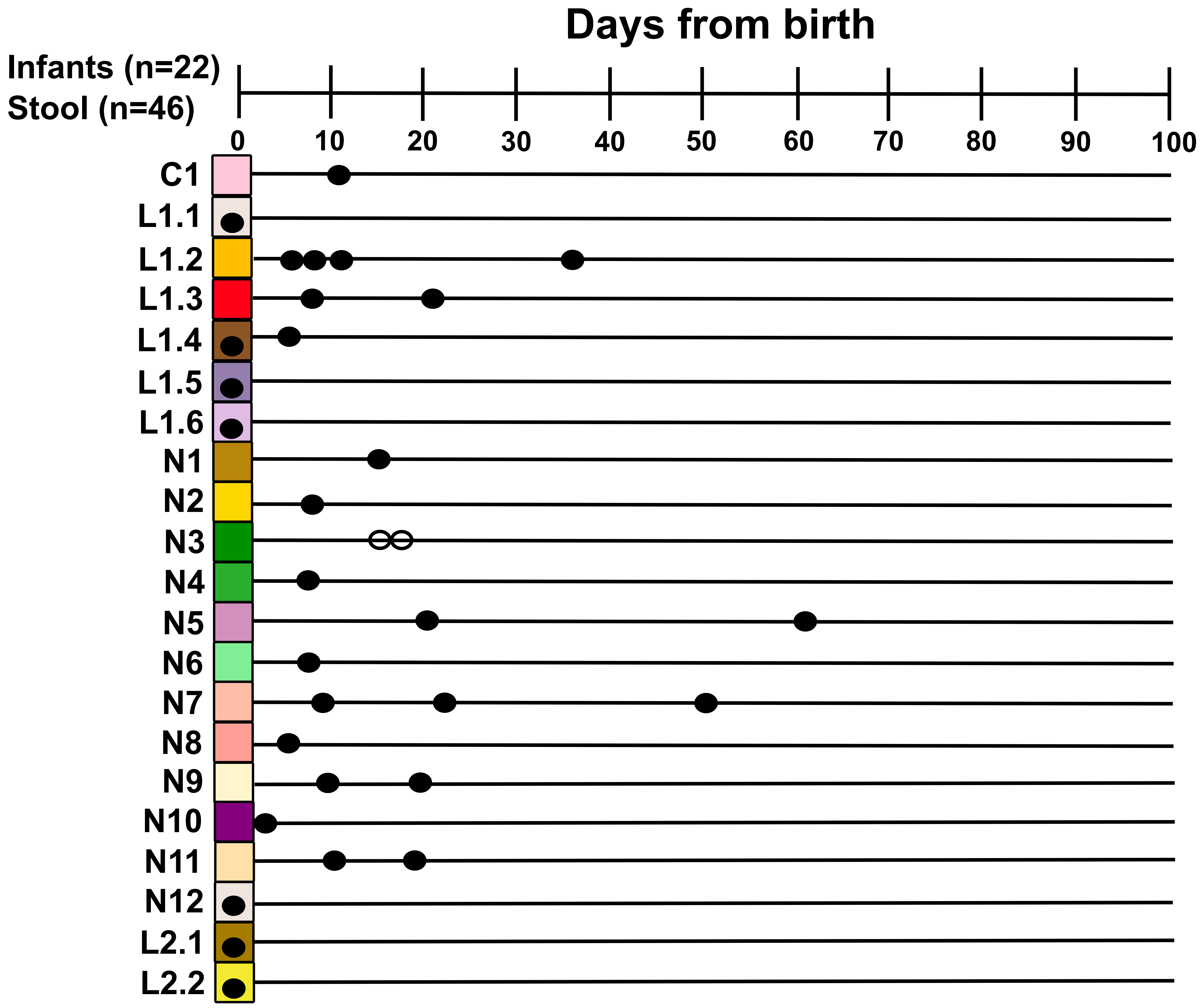

### Fig. S2

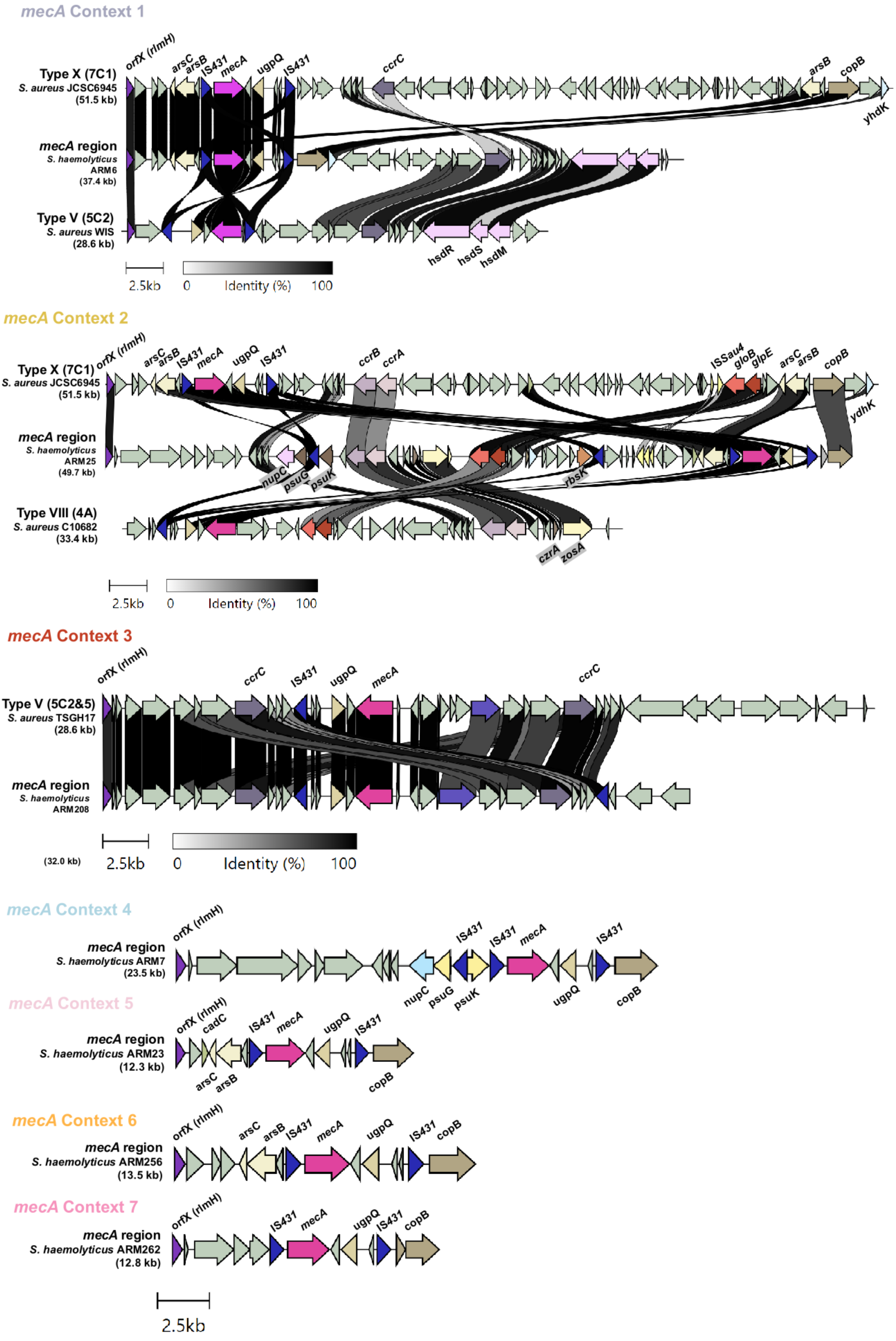

### Fig. S3

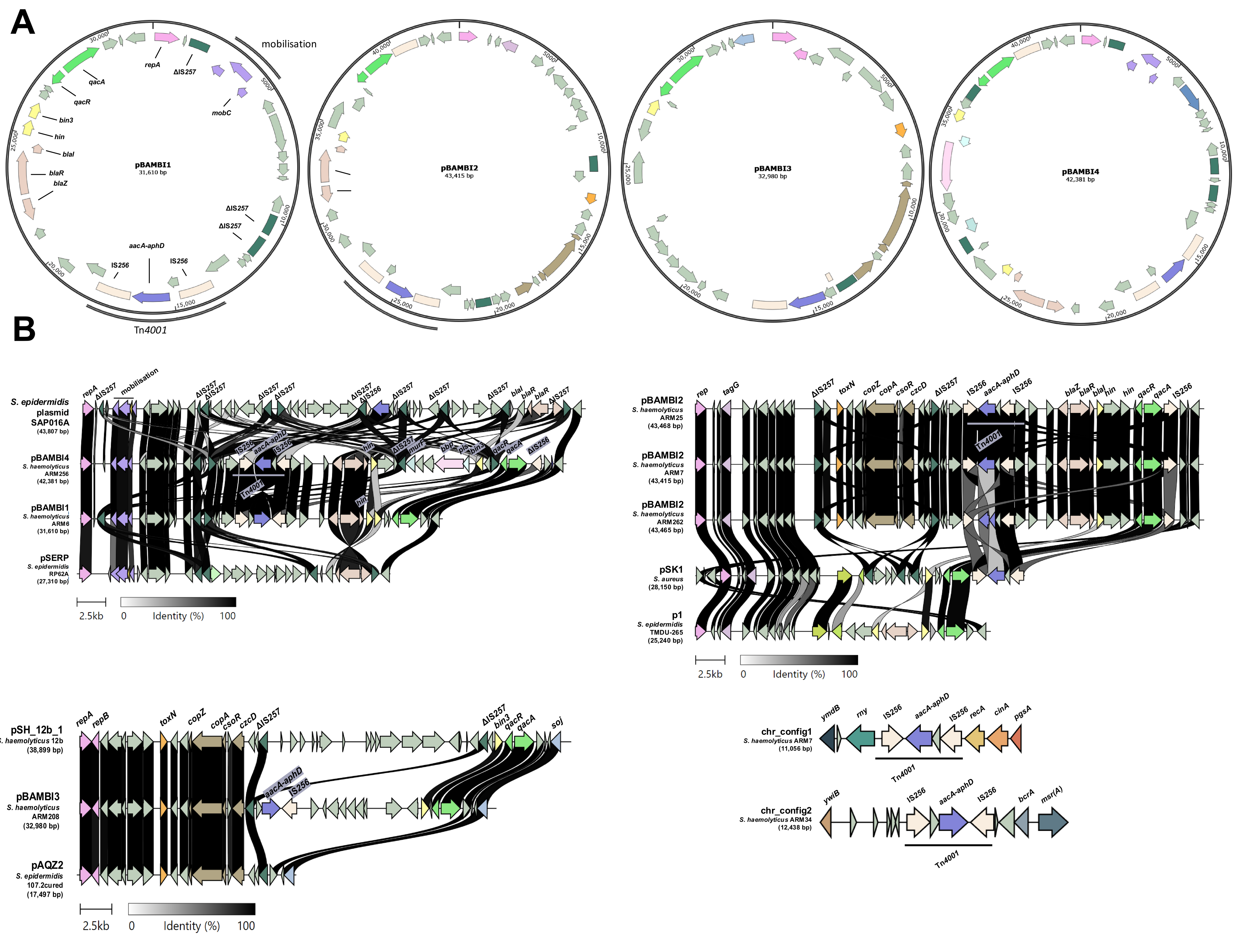
