## Supplementary material for "*Staphylococcus haemolyticus* is a reservoir of antibiotic resistance genes in the preterm infant gut": Table S1

**Supplementary Table 1.** Reference dataset of SCC*mec*-encoded elements.

| SCC*mec* element | Source | Genbank Accession |
| --- | --- | --- |
| orfX | *Staphylococcus aureus* DNA, type-I staphylococccal cassette chromosome mec: strain NCTC10442 | AB033763.2 |
| mecA | Staphylococcus aureus DNA, type-I staphylococccal cassette chromosome mec: strain NCTC10442 | AB033763.2 |
| mecR1 | Staphylococcus aureus DNA, type-I staphylococccal cassette chromosome mec: strain NCTC10442 | AB033763.2 |
| IS431 | Staphylococcus aureus DNA, type-I staphylococccal cassette chromosome mec: strain NCTC10442 | AB033763.2 |
| mecI | Staphylococcus aureus DNA, type-II staphylococcal cassette chromosome mec: strain N315 | D86934.2 |
| ccrA2 | Staphylococcus aureus DNA, type IVc staphylococcal cassette chromosome mec, strain:80s-2 | AB245471.1 |
| ccrA3 | Staphylococcus aureus gene for cassette chromosome recombinase A, cassette chromosome recombinase B, complete cds, strain 85/1340 | AB014437.1 |
| ccrA3 | Staphylococcus aureus gene for cassette chromosome recombinase A, cassette chromosome recombinase B, complete cds, strain 85/2082 | AB014436.1 |
| ccrA3 | Staphylococcus aureus DNA, left extremity of Type-III SCCmec (Staphylococcal cassette chromosome mec) and it's flanking chromosomal region, strain:85/3907 | AB047088.2 |
| ccrA3 | Staphylococcus aureus isolate A-44K9S cassette chromosome recombinase A (ccrA3) and cassette chromosome recombinase B (ccrB3) genes, complete cds | DQ196432.1 |
| ccrA2 | Staphylococcus epidermidis isolate A-53DS cassette chromosome recombinase A (ccrA2) and cassette chromosome recombinase B (ccrB2) genes, complete cds, | DQ196433.1 |
| ccrA2 | Staphylococcus aureus DNA, type IVE Staphylococcal Cassette Chromosome mec (SCCmec), strain AR43/3330.1 | AJ810121.1 |
| ccrA2 | Staphylococcus aureus strain C98-370 CcrA and CcrB genes, complete cds | DQ514327.1 |
| ccrA2 | Staphylococcus aureus strain C99-529 CcrB and CcrA genes, complete cds | DQ514328.1 |
| ccrA4 | Staphylococcus aureus strain CHE482 cassette chromosome recombinase A (ccrA4-1) and cassette chromosome recombinase B (ccrB4-1) genes, complete cds | EF126185.1 |
| ccrA4 | Staphylococcus aureus strain CHE482 cassette chromosome recombinase A (ccrA4-2) and cassette chromosome recombinase B (ccrB4-2) genes, complete cds | EF126186.1 |
| ccrA2 | Staphylococcus epidermidis isolate CS8 cassette chromosome recombinase A (ccrA2) and cassette chromosome recombinase B (ccrB2) genes, complete cds | DQ196434.1 |
| ccrA4 | Staphylococcus epidermidis ATCC 12228, complete sequence | NC_004461.1 |
| ccrA2 | Staphylococcus epidermidis ATCC 12228, complete sequence | NC_004461.1 |
| ccrA2 | Staphylococcus warneri cassette chromosome recombinase A (ccrA2) and cassette chromosome recombinase B (ccrB2) genes, complete cds, | DQ225180.1 |
| ccrA1 | Staphylococcus aureus subsp. aureus COL, complete sequence | NC_002951.2 |
| ccrA2 | Staphylococcus warneri cassette chromosome recombinase A (ccrA2) and cassette chromosome recombinase B (ccrB2) genes, complete cds | DQ225180.1 |
| ccrA1 | Staphylococcus hominis DNA, staphylococcal cassette chromosome : strain GIFU12263 | AB063171.1 |
| ccrA4 | Staphylococcus aureus strain HDE288 type-VI SCCmec element, complete sequence | AF411935.3 |
| ccrA2 | Staphylococcus aureus strain J28 CcrA and CcrB genes, complete cds | DQ514329.1 |
| ccrA2 | Staphylococcus aureus strain J35 CcrB and CcrA genes, complete cds | DQ514330.1 |
| ccrA2 | Staphylococcus aureus strain J52 CcrA and CcrB genes, complete cds | DQ514331.1 |
| ccrA2 | Staphylococcus aureus DNA, type-IV.1 (IVa) staphylococcal cassette chromosome mec: strain CA05(JCSC1968) | AB063172.2 |
| ccrA2 | Staphylococcus aureus DNA, type-IV.2 (IVb) staphylococcal cassette chromosome mec: strain JCSC1978(8/6-3P) | AB063173.1 |
| ccrA2 | Staphylococcus aureus DNA, type-IIb staphylococcal cassette chromosome mec: strain JCSC3063, | AB127982.1 |
| ccrA2 | Staphylococcus aureus gene for cassette chromosome recombinase A, cassette chromosome recombinase B, complete cds, strain 85/1340 | AB014437.1 |
| ccrA2 | Staphylococcus aureus DNA, type-IV.1 (IVa) staphylococcal cassette chromosome mec: strain JCSC4744 | AB266531.1 |
| ccrA2 | Staphylococcus aureus DNA, type-IV.3 (IVc) staphylococcal cassette chromosome mec:strain JCSC4788 | AB266532.1 |
| ccrA3 | Staphylococcus pseudintermedius SCCmec cassette, strain KM1381 | AM904732.1) |
| ccrA5 | Staphylococcus pseudintermedius SCCmec cassette, strain KM241 | AM904731.1 |
| ccrA2 | Staphylococcus aureus subsp. aureus JH9, complete sequence | NC_009487.1 |
| ccrA2 | Staphylococcus aureus strain M03-68 type-IVg SCCmec element, complete sequence | DQ106887.1 |
| ccrA2 | Staphylococcus aureus DNA, type-II staphylococcal cassette chromosome mec: strain N315 | D86934.2 |
| ccrA1 | Staphylococcus aureus DNA, type-I staphylococccal cassette chromosome mec: strain NCTC10442 | AB033763.2 |
| ccrA2 | Staphylococcus aureus subsp. aureus MRSA252, complete sequence | NC_002952.2 |
| ccrA2 | Staphylococcus aureus DNA, type IVc staphylococcal cassette chromosome mec, strain:NN1 | AB245470.1 |
| ccrA1 | Staphylococcus aureus subsp. aureus MSSA476, complete sequence | NC_002953.3 |
| ccrA2 | Staphylococcus aureus DNA, J1 region of typeII.4 staphylococcal cassette chromosome mec, strain RN7170 | AB261975.1 |
| ccrA2 | Staphylococcus epidermidis strain SE5 CcrA and CcrB genes, complete cds | DQ514332.1 |
| ccrA2 | Staphylococcus aureus subsp. aureus Mu3, complete sequence | NC_009782.1 |
| ccrA2 | Staphylococcus epidermidis strain SE50 CcrA and CcrB genes, complete cds | DQ514334.1 |
| ccrA2 | Staphylococcus epidermidis strain SE63 CcrA and CcrB genes, complete cds | DQ514335.1 |
| ccrA2 | Staphylococcus aureus subsp. aureus Mu50, complete sequence | NC_002758.2 |
| ccrA2 | Staphylococcus epidermidis strain SE7 CcrA and CcrB genes, complete cds | DQ514333.1 |
| ccrA1 | Staphylococcus saprophyticus TSU 33 DNA, Staphylococcal Cassette Chromosome mec | AB353724.1 |
| ccrA2 | Staphylococcus aureus subsp. aureus MW2, complete sequence | NC_003923.1 |
| ccrB2 | Staphylococcus aureus SCCmec type IVc element, complete sequence | AY271717.1 |
| ccrA2 | Staphylococcus epidermidis RP62A, complete sequence | NC_002976.3 |
| ccrB2 | Staphylococcus aureus DNA, type IVc staphylococcal cassette chromosome mec, strain:80s-2 | AB245471.1 |
| ccrB3 | Staphylococcus aureus gene for cassette chromosome recombinase A, cassette chromosome recombinase B, complete cds, strain 85/1340 | AB014437.1 |
| ccrA1 | Staphylococcus saprophyticus subsp. saprophyticus ATCC 15305 = NCTC 7292, complete sequence | NC_007350.1 |
| ccrB3 | Staphylococcus aureus gene for cassette chromosome recombinase A, cassette chromosome recombinase B, complete cds, strain 85/2082 | AB014436.1 |
| ccrB3 | Staphylococcus aureus DNA, left extremity of Type-III SCCmec (Staphylococcal cassette chromosome mec) and it's flanking chromosomal region, strain:85/3907 | AB047088.2 |
| ccrA2 | Staphylococcus aureus subsp. aureus USA300_FPR3757, complete sequence | NC_007793.1 |
| ccrA2 | Staphylococcus aureus subsp. aureus USA300_TCH1516, complete sequence | NC_010079.1 |
| ccrB3 | Staphylococcus aureus isolate A-44K9S cassette chromosome recombinase A (ccrA3) and cassette chromosome recombinase B (ccrB3) genes, complete cds | DQ196432.1 |
| ccrB2 | Staphylococcus epidermidis isolate A-53DS cassette chromosome recombinase A (ccrA2) and cassette chromosome recombinase B (ccrB2) genes, complete cds | DQ196433.1 |
| ccrB2 | Staphylococcus aureus DNA, type IIE Staphylococcal Cassette Chromosome mec (SCCmec), strain AR13.1/3330.2 | AJ810120.1 |
| ccrB2 | Staphylococcus aureus DNA, type IVE Staphylococcal Cassette Chromosome mec (SCCmec), strain AR43/3330.1 | AJ810121.1 |
| ccrB2 | Staphylococcus aureus strain C98-370 CcrA and CcrB genes, complete cds | DQ514327.1 |
| ccrB2 | Staphylococcus aureus strain C99-529 CcrB and CcrA genes, complete cds | DQ514328.1 |
| ccrB4 | Staphylococcus aureus strain CHE482 cassette chromosome recombinase A (ccrA4-1) and cassette chromosome recombinase B (ccrB4-1) genes, complete cds | EF126185.1 |
| ccrB4 | Staphylococcus aureus strain CHE482 cassette chromosome recombinase A (ccrA4-2) and cassette chromosome recombinase B (ccrB4-2) genes, complete cds | EF126186.1 |
| ccrB2 | Staphylococcus epidermidis ATCC 12228, complete sequence | NC_004461.1 |
| ccrB4 | Staphylococcus epidermidis ATCC 12228, complete sequence | NC_004461.1 |
| ccrB2 | Staphylococcus epidermidis isolate CS8 cassette chromosome recombinase A (ccrA2) and cassette chromosome recombinase B (ccrB2) genes, complete cds | DQ196434.1 |
| ccrB2 | Staphylococcus warneri cassette chromosome recombinase A (ccrA2) and cassette chromosome recombinase B (ccrB2) genes, complete cds | DQ225180.1 |
| ccrB1 | Staphylococcus hominis DNA, staphylococcal cassette chromosome : strain GIFU12263 | AB063171.1 |
| ccrB1 | Staphylococcus aureus subsp. aureus COL, complete sequence | NC_002951.2 |
| ccrB1 | Staphylococcus aureus strain HDE288 type-VI SCCmec element, complete sequence | AF411935.3 |
| ccrB2 | Staphylococcus aureus strain J28 CcrA and CcrB genes, complete cds | DQ514329.1 |
| ccrB2 | Staphylococcus aureus strain J35 CcrB and CcrA genes, complete cds | DQ514330.1 |
| ccrB2 | Staphylococcus aureus strain J52 CcrA and CcrB genes, complete cds | DQ514331.1 |
| ccrB2 | Staphylococcus aureus DNA, type-IV.1 (IVa) staphylococcal cassette chromosome mec: strain CA05(JCSC1968) | AB063172.2 |
| ccrB2 | Staphylococcus aureus DNA, type-IV.2 (IVb) staphylococcal cassette chromosome mec: strain JCSC1978(8/6-3P) | AB063173.1 |
| ccrB2 | Staphylococcus aureus DNA, type-IIb staphylococcal cassette chromosome mec: strain JCSC3063 | AB127982.1 |
| ccrB2 | Staphylococcus aureus gene for cassette chromosome recombinase A, cassette chromosome recombinase B, complete cds, strain 85/1340 | AB014437.1 |
| ccrB2 | Staphylococcus aureus DNA, type-IV.1 (IVa) staphylococcal cassette chromosome mec: strain JCSC4744 | AB266531.1 |
| ccrB2 | Staphylococcus aureus DNA, type-IV.3 (IVc) staphylococcal cassette chromosome mec:strain JCSC4788 | AB266532.1 |
| ccrB3 | Staphylococcus pseudintermedius SCCmec cassette, strain KM1381 | AM904732.1 |
| ccrB3 | Staphylococcus pseudintermedius SCCmec cassette, strain KM241 | AM904731.1 |
| ccrB2 | Staphylococcus aureus strain M03-68 type-IVg SCCmec element, complete sequence | DQ106887.1 |
| ccrB2 | Staphylococcus aureus subsp. aureus JH9, complete sequence | NC_009487.1 |
| ccrB1 | Staphylococcus aureus DNA, type-I staphylococccal cassette chromosome mec: strain NCTC10442 | AB033763.2 |
| ccrB2 | Staphylococcus aureus DNA, type IVc staphylococcal cassette chromosome mec, strain:NN1 | AB245470.1 |
| ccrB2 | Staphylococcus aureus subsp. aureus MRSA252, complete sequence | NC_002952.2 |
| ccrB1 | Staphylococcus aureus subsp. aureus MSSA476, complete sequence | NC_002953.3 |
| ccrB2 | Staphylococcus aureus DNA, J1 region of typeII.4 staphylococcal cassette chromosome mec, strain RN7170 | AB261975.1 |
| ccrB2 | Staphylococcus epidermidis strain SE5 CcrA and CcrB genes, complete cds | DQ514332.1 |
| ccrB2 | Staphylococcus aureus subsp. aureus Mu3, complete sequence | NC_009782.1 |
| ccrB2 | Staphylococcus epidermidis strain SE50 CcrA and CcrB genes, complete cds | DQ514334.1 |
| ccrB2 | Staphylococcus epidermidis strain SE63 CcrA and CcrB genes, complete cds | DQ514335.1 |
| ccrB2 | Staphylococcus epidermidis strain SE7 CcrA and CcrB genes, complete cds | DQ514333.1 |
| ccrB2 | Staphylococcus aureus subsp. aureus Mu50, complete sequence | NC_002758.2 |
| ccrB7 (Genbank Acc:) | Staphylococcus saprophyticus TSU 33 DNA, Staphylococcal Cassette Chromosome mec | AB353724.1 |
| ccrC1 | Staphylococcus aureus DNA, type-V staphylococcal cassette chromosome mec: strain JCSC3624(WIS) | AB121219.1 |
| ccrB2 | Staphylococcus aureus subsp. aureus MW2, complete sequence | NC_003923.1 |
| ccrC2 | Staphylococcus aureus hypothetical protein gene, partial cds; and hypothetical proteins, cassette chromosome recombinase C (ccrC), and hypothetical proteins genes, complete cds, | AY894416.1 |
| ccrB2 | Staphylococcus epidermidis RP62A, complete sequence | NC_002976.3 |
| ccrC5 | Staphylococcus haemolyticus JCSC1435 DNA, complete genome | AP006716.1 |
| ccrC6 | Staphylococcus haemolyticus strain 25-60 cassette chromosome recombinase C6 (ccrC6) gene, complete cds | EF190467.1 |
| ccrB2 | Staphylococcus aureus subsp. aureus USA300_FPR3757, complete sequence | NC_007793.1 |
| ccrC7 | Staphylococcus epidermidis strain 13-48 cassette chromosome recombinase C7 (ccrC7) gene, complete cds | EF190468.1 |
| ccrC8 | Staphylococcus aureus DNA, type VII staphylococcal cassette chromosome mec, staphylococcal orfX-orfY region carrying SCCmec VII, strain: PM1 | AB462393.1 |
| ccrB2 | Staphylococcus aureus subsp. aureus USA300_TCH1516, complete sequence | NC_010079.1 |
| ccrC3 | Staphylococcus aureus DNA, type-III staphylococcal cassette chromosome mec and SCCmercury: strain 85/2082 | AB037671.1 |
| ccrC4 | Staphylococcus aureus M type 1 capsular polysaccharide biosynthesis gene cluster, complete sequence and unknown genes | U10927.2 |
| ccrC9 | Staphylococcus saprophyticus subsp. saprophyticus ATCC 15305 = NCTC 7292, complete sequence | NC_007350.1 |
